# The Fly Disco: Hardware and software for optogenetics and fine-grained fly behavior analysis

**DOI:** 10.1101/2024.11.04.621948

**Authors:** Alice A. Robie, Adam L. Taylor, Catherine E. Schretter, Mayank Kabra, Kristin Branson

## Abstract

In the fruit fly, *Drosophila melanogaster*, connectome data and genetic tools provide a unique opportunity to study complex behaviors including navigation, mating, aggression, and grooming in an organism with a tractable nervous system of 140,000 neurons. Here we present the Fly Disco, a flexible system for high quality video collection, optogenetic manipulation, and fine-grained behavioral analysis of freely walking and socializing fruit fly groups. The data collection hardware and software automates the collection of videos synced to programmable optogenetic stimuli. Key pipeline features include behavioral analysis based on trajectories of 21 keypoints and optogenetic-specific summary statistics and data visualization. We created the *multifly* dataset for pose estimation that includes 9701 examples enriched in complex behaviors. All hardware designs, software, and the *multifly* dataset are freely available.

## Introduction

Neuroscience in the model organism, *Drosophila melanogaster*, has undergone a paradigm shift as connectomic projects catalog the neurons and circuits of the central nervous system (***Scheffer et al., 2020***; ***Buhmann et al., 2021***; ***Devineni, 2024***; ***Dorkenwald et al., 2024***). From these wiring diagrams, computational analysis can generate new hypotheses of circuit function (***Shiu et al., 2023***; ***Hulse et al., 2021***). Directly testing such hypotheses requires manipulating neural activity in behaving flies. Genetic tools for spatially-targeted, temporally-restricted manipulations of neurons have been developed and tested in flies (***Klapoetke et al., 2014***; ***Mauss et al., 2017***; ***Emiliani et al., 2022***; ***Meissner et al., 2024***; ***Luan et al., 2020***). Previous studies have demostarted the utility of these genetic tools for activation and inactivation of individual cell types for functional investigation of sensory neurons (***Wu et al., 2016***), descending interneurons (***Namiki et al., 2018***; ***Cande et al., 2018***), and the courtship song circuit (***Lillvis et al., 2024***) in single or paired fly assays. There exists a need for a flexible system for recording, manipulating, and analyzing a large variety of behaviors (including locomotion and social interactions) in a detailed quantitative manner.

Here, we present the Fly Disco, a system from data collection through fine-grained behavioral analysis for performing high-throughput optogenetic behavioral manipulations in groups of freely walking fruit flies. This system was designed to produce video for which automated machine vision tools can be used quantify how flies’ use their limbs and wings during behaviors such as walking, grooming, courtship, and aggression. We engineered data collection hardware and software to maximize throughput while minimizing human error. The flies are recorded from above in a 50 mm diameter, 3.5 mm tall chamber with a sloped ceiling. The chamber is illuminated from below with a programmable multicolor LED array that provides optogenetic stimulation in three colors and infrared (IR) light for imaging the flies. The data collection software automates video acquisition, digital metadata collection, and optogenetic stimulus generation. We developed a modular analysis pipeline that performs body tracking, sex-classification, keypoint tracking, behavior classification, and outputs stimulus-trigger behavioral quantification of the effects of neural manipulations. We provide the all the design files, parts lists needed to build the rig, and the software for data acquisition and the analysis pipeline.

Training a machine learning-based pose tracker of sufficient accuracy to study complex motor and social behaviors in groups of flies such as grooming, courtship and aggression requires a high quality training dataset containing a large number of annotated examples of flies performing these behaviors. These are inherently more difficult examples for both humans and machine learning, as they involve occlusions, variable body poses, and identity assignment. Our *multifly* dataset, consisting of 9701 examples, was constructed to include many of these difficult examples. Trained annotators labeled every example with all 21 keypoints, determining the location of even occluded keypoints by using temporal context and domain knowledge. Hard examples were added to the training set iteratively with annotators finding examples with errors in tracking prediction and adding those to the training dataset. The test set was designed to quantify the generalization of the tracker on novel data and the effect of fly behavior on prediction accuracy.

Our keypoint tracker was accurate with a mean error of .04 mm, or 1.5% of a body length of a 2.7 mm *Drosophila melanogaster*, on test data enriched with difficult examples. We demonstrated the utility of our accurate pose tracking in two ways. First, we trained classifiers for grooming and social interaction behaviors from keypoint trajectories. Analysis of the leg tip positions relative to a target fly revealed patterns in how flies target their legs during courtship and aggression behaviors. Second, we created a analysis module to quantified the legs movements in groups of freely walking flies (***Mendes et al., 2013***; ***Wosnitza et al., 2013***; ***DeAngelis et al., 2019***).

## Results

### Hardware and Data Collection Software

We developed an assay designed for machine-vision and -learning analysis. We video-recorded groups of 10 freely walking flies in the 50mm diameter, 3.5mm tall chamber from above at 1MP and 150 fps (***Figure 1***). We designed the transparent lid of chamber to reduce walking on the ceiling and walls and named it the *bubble* due to its flattened dome shape (***Simon and Dickinson, 2010***; ***Berman et al., 2014***). Restricting the flies to the backlit floor of the chamber, a plane orthogonal to the camera, provided a consistent, high-contrast image of the body, legs, and wings that facilitated high-quality pose tracking. The recording chamber was assembled by sliding the bubble into place above the transparent floor of the cartridge. To enable high-throughput data collection, this design allowed quick loading and unloading of flies, reproducible placement within the recording apparatus, and easy disassembly for cleaning between videos. The black insert that holds the bubble created a clear demarcation of the chamber boundary facilitating automated arena detection in the analysis pipeline (see Fly Bubble behavioral chamber).

**Figure 1.**
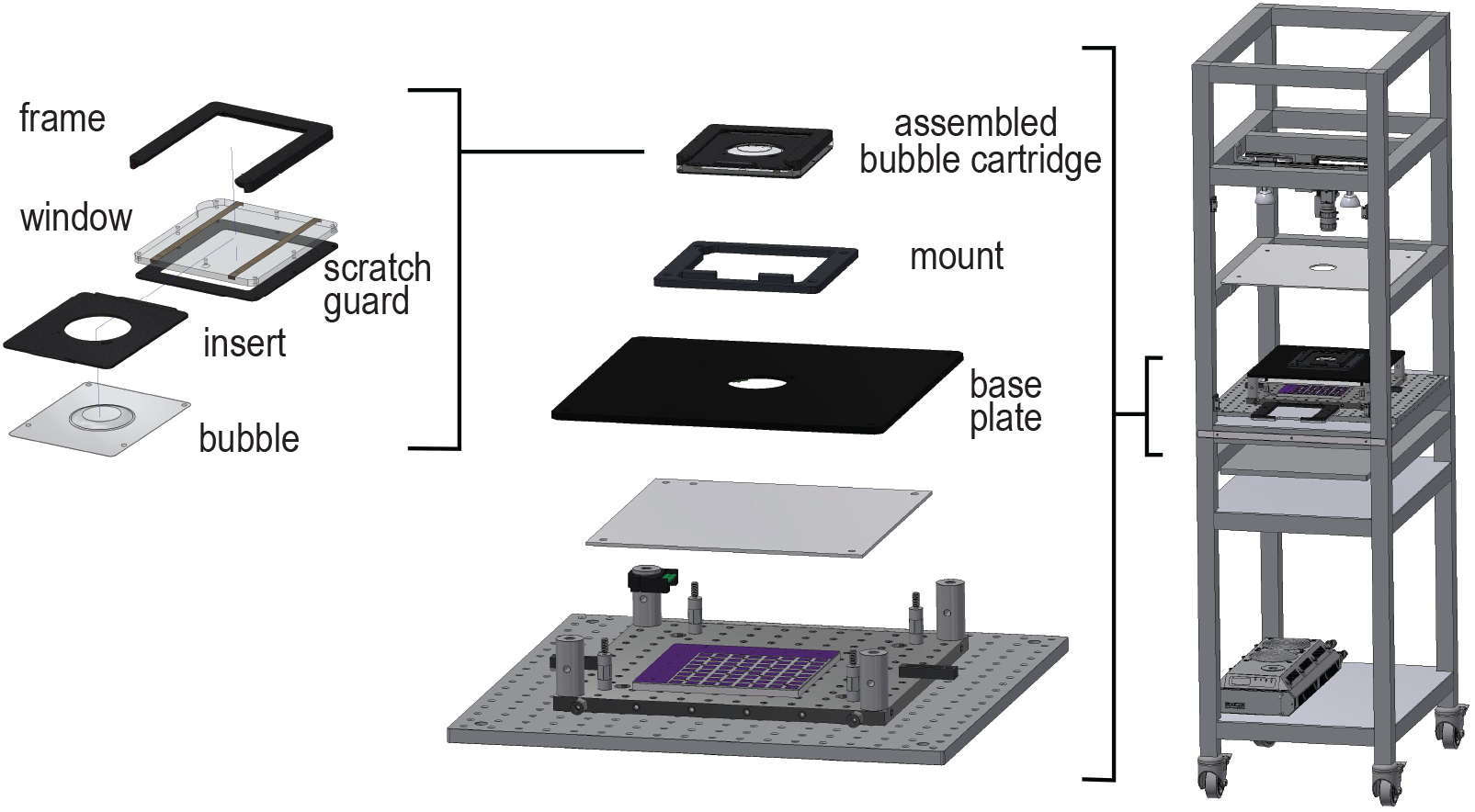
Schematic of Fly Disco Hardware. The left-hand panel displays the individual components of the Fly Bubble cartridge: The frame, window and scratch guard are screwed together to form the empty cartridge into which slides the bubble held by the insert to form the recording chamber. The middle panel shows an exploded view of the rig components. The LED array is thermally glued to a metal heat-sink and attached to the water-cooled breadboard. Support posts for the LED array’s diffuser (innner posts) and the metal base plate (outer posts) are also secured to the water-cooled breadboard. The indicator LED electronics are attached at the top of the rear-left post, and are electrically connected via spring-loaded contacts to the the indicator LEDs (not illustrated) that are screwed to the bottom of the base plate. The asymmetrical 3D-printed mounting frame for the bubble cartridge is secured to the baseplate, and the assembled bubble cartridge is placed in the mount for recording. The right-hand panel is schematic of the assembled Fly Disco Rig on its cart including the loading frame, the mounted camera, the visible lights and their diffuser, and the CPU radiator for the water-cooled breadboard. Not pictured are the blackout panels, tubing, wires, the computer, and the power supplies.

The bubble chamber is illuminated from below with a custom high-intensity LED array that generates programmable, multi-color optogenetic stimuli and provides constant IR light for imaging (***Figure 2***A). To provide even, multi-color illumination across the chamber, compact groups of color and IR LEDs tile the board. We measured the power-intensity curves of LEDs within the behavioral chamber (***Figure 2***B). The LEDs generate sufficient light intensity for optogenetic activation with red-light gated cation channel (CsChR) (***Klapoetke et al., 2014***) and inactivation with green-light gated anion channel (GtACR1) (***Mauss et al., 2017***). An IR-pass filter on the camera blocks the optogenetic stimuli, preventing it from interferring with fly imaging. To record the activity of the multicolor LEDs, the board uses three IR LEDs in the camera’s field-of-view, positioned outside the arena, to reproduce the optogenetic stimuli pattern. This provides a reliable method of synchronizing the optogenetic stimulation to the video of fly behavior. Taken together, the Fly Bubble chamber and the custom multi-color LED array form the core of the Fly Disco hardware. Further details of the rig are described in Fly Disco Hardware.

**Figure 2.**
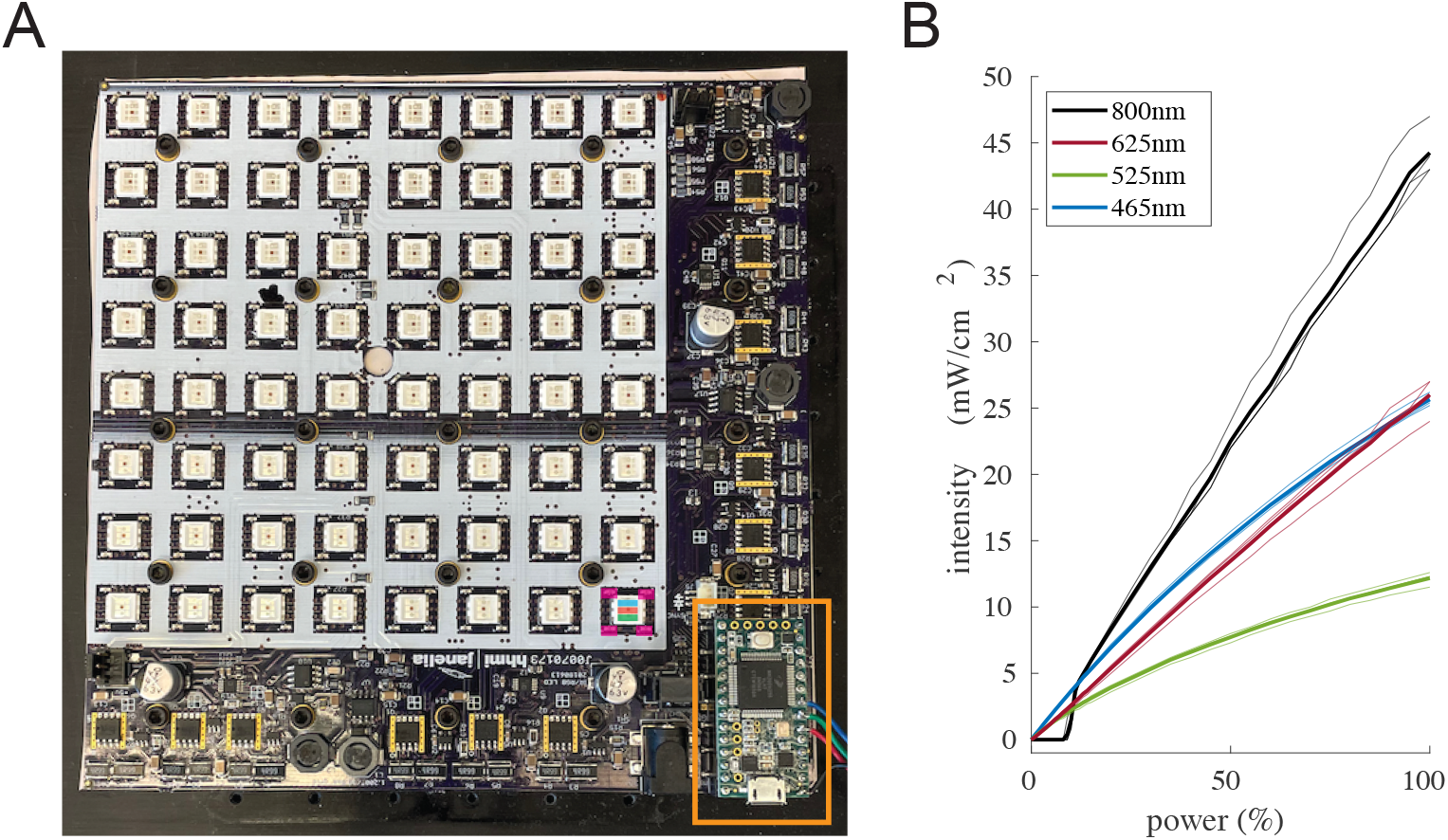
(A) Photograph of the LED board. The LED array (white background) is tiled with groups of IR and RGB LEDs. In the lower left corner, a single group is highlighted: 4 IR LEDs (in magenta) around a central tricolor LED (in green, red, and blue). The Arduino^®^ control board is indicated by the orange box. (B) The power input vs intensity output is plotted for the 4 LED colors IR (800nm), red (625nm), green (525nm) and blue (465nm) as measured with a Thor Labs PM100D power meter.

A Flea3 USB 3.0 camera mounted above the recording chamber collects video via our custom data acquisition software, Fly Bowl Data Capture (FBDC) (***Robie et al., 2017***). The software automates data collection and recording digital metadata for each video. It is designed to minimize human error and maximize the data collection rate (***Figure 3***). We modified to the software to support USB 3.0 cameras and optogenetic experiments through the control of the programmable multi-color LED array. For USB 3.0 cameras, FBDC does not interface with the camera directly, instead it controls video acquisition through our BIAS camera software. FBDC functionality is detailed in Data acquisition software.

**Figure 3.**
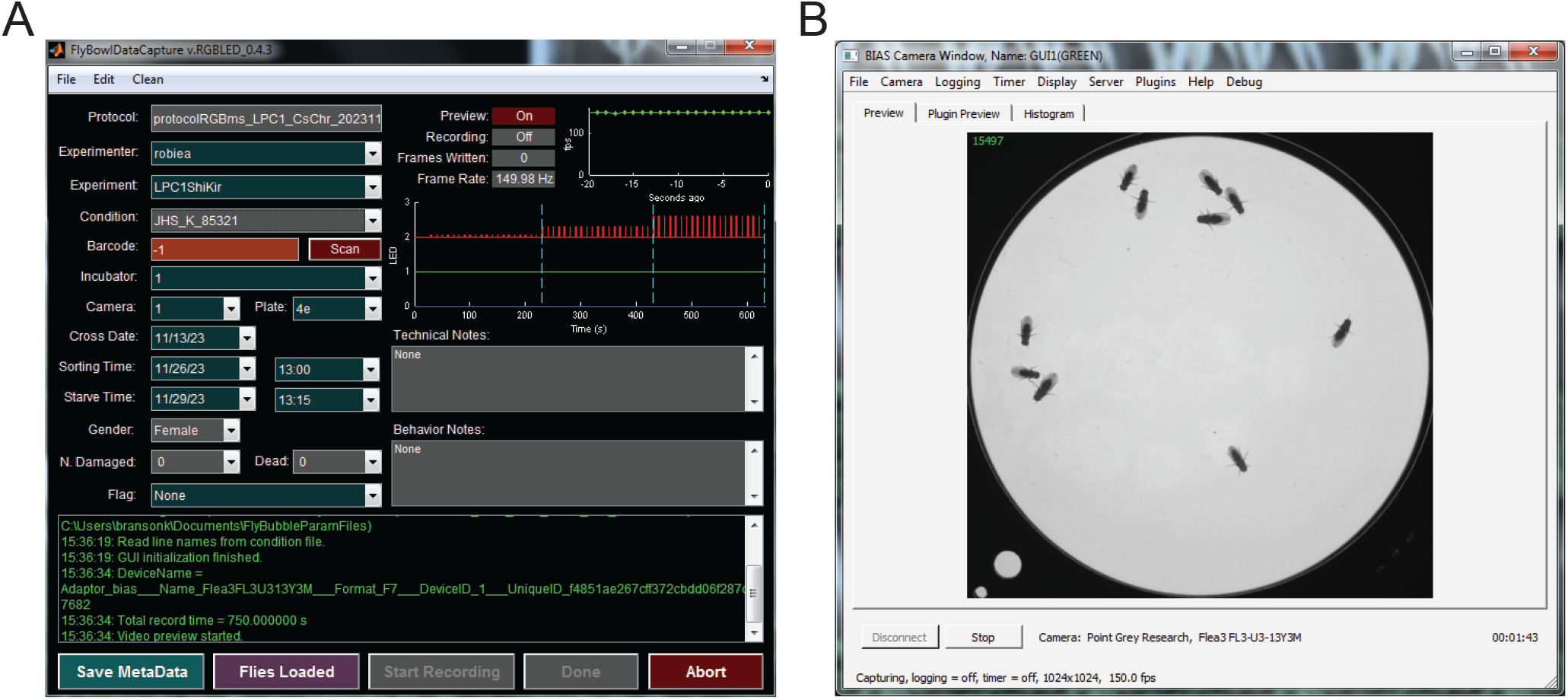
Data collection software (A) The FBDC GUI displays experimental metadata in the upper-left. Real-time status of the recording, FPS, and progress through the LED stimuli are shown in the upper-right, and below this free-form text boxes for notes that are saved in the digital metadata. Real-time experimental software outputs are shown in green, and the action button for running the experiment are the bottom row of the GUI. (B) The BIAS window showing a video preview of flies in the Fly Bubble chamber. Video acquisition is automated via FBDC.

### Data Analysis Pipeline

Our Fly Disco apparatus allows high-throughput collection of large amounts of video of fly behaviors. With four Fly Disco rigs operated in parallel by one researcher, we collected one 6.5 minute long videos every 3.25 minutes, 18 videos/hour. At 150 fps, this results in ∼1 million frames of video data per hour of data collection. To efficiently analyze the flies’ behavior in these videos, we created a modular pipeline, the Fly Disco Analysis Pipeline, that automates multiple stages of data processing (***Figure 4***, Fly Disco Analysis Pipeline). The final output of the pipeline is a fine-grained, quantitative description of changes in fly behavior due to experimental manipulation. This pipeline is a major revision of our previously published pipeline (***Robie et al., 2017***) with novel functionality added to process optogenetic experiments and perform keypoint tracking, and substantial updates to the majority of the stages (***Figure 4***). However, the general framework of data processing remains intact: modular stages are run sequentially and governed by user defined configuration files for each stage (***Figure 5, Table 2***). This framework gives the pipeline the flexibility to process videos collected from different apparatuses, of varying lengths, with and without optogenetic stimulation, and with different numbers or sexes of flies being recorded. Our analysis includes machine learning-based models trained on data from our Fly Disco rigs. Adaptation of our pipeline to a new assays will either require videos that are qualitatively similar to ours, or fine-tuning of the ML models to the new types of data.

**Figure 4.**
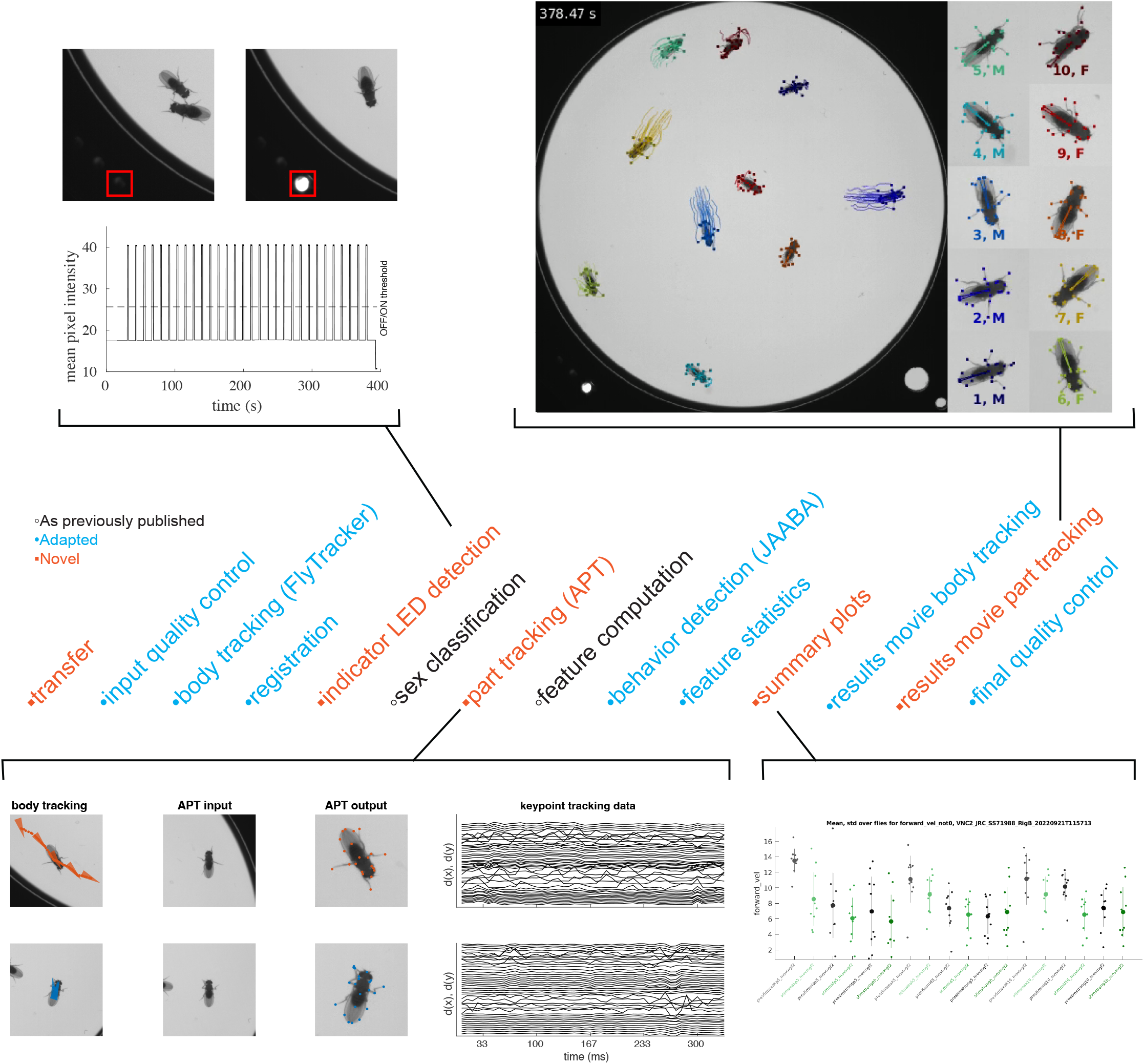
Fly Disco Analysis pipeline. The stages of the pipeline are shown in the center of the figure. This pipeline builds on our previous data analysis pipeline ***Robie et al. (2017***), and each stage is color based on novel (orange), modified (blue), and unmodified (black). The four novel data processing stages are illustrated. *indicator LED detection* detects the status of indicator LED in each frame to digitize the optogenetic stimuli timeline: when an IR LED is illuminated, the corresponding color of the LED array is active. *Part tracking (APT)* detects 21 keypoints on the fly’s body in each frame producing movement information for the head, thorax, abdomen and each limb. *summary plots* produce statistics summarizing the flies’ behavior such as mean speed of the flies during the first second of each optogenetic stimulation period. The computed metrics can be customized using a controlled vocabulary. *results movie part tracking* creates a short video of user-defined snippets showing the output of keypoint tracking.

**Figure 5.**
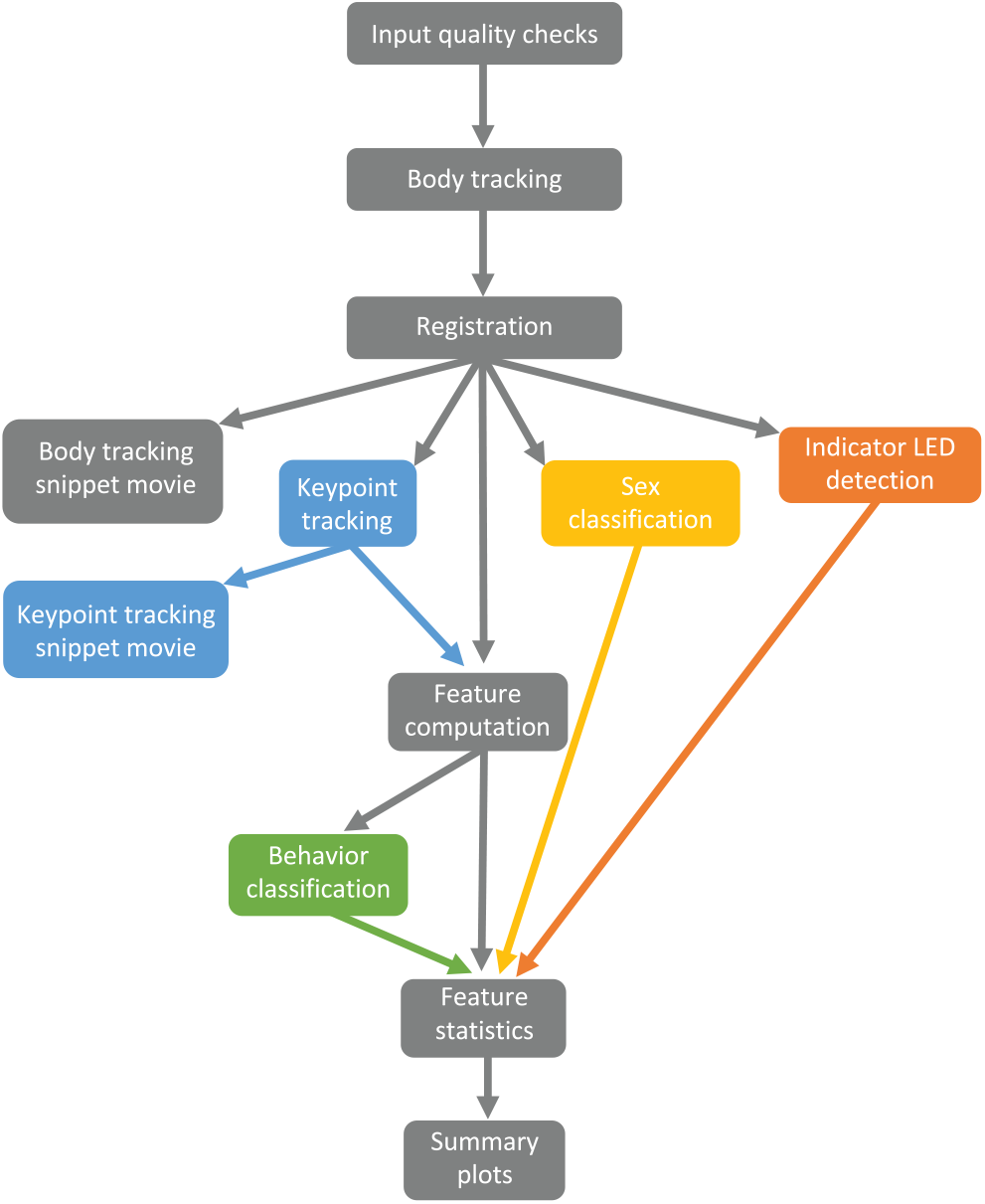
FDA data processing flowchart. This flowchart shows how data moves through the stages of the FDA with each arrow representing required pre-processing for the current stage. Grey boxes represent the basic version of the pipeline and each color represents optional additional stages of processing. The complete quality checks (not illustrated) require inputs from every active stage of the pipeline.

For each fly in each video, we automatically tracked the body position and orientation, classified the sex, and predicted the location of 21 keypoints on the fly (Multifly Pose Tracker: Dataset and Performance Section). From this trajectory data, we computed engineered behavioral metrics using short temporal contexts such as speed, relative position to other flies, and limb movement relative the body or another limb. These behavior metrics, termed *per-frame features*, describe the behavioral state of the fly in a given frame. These low-level behavioral metrics can be summarized to described the animal’s behavior throughout the movie such as mean turning rate. To quantify more complex behaviors such as grooming or social interactions, we used our behavior classification software, JAABA, which uses supervised machine-learning to produce behavior classifiers based on the per-frame features (***Kabra et al. (2013***), Behavior classification: JAABA). The pipeline uses these pretrained classifiers to predict the behavior of each fly in every frame of the video.

For analysis of optogenetic experiments, we added stimulus-triggered averaging to quantify the behavioral effects of neuronal manipulation. Different optogenetic experiments utilize various stimulation protocols, often involving repeated presentation of the stimuli at different intensity levels. In order to make our automated analysis flexible, we developed a controlled vocabulary from which users can construct analysis windows over which summary statistics are calculated. For example, suppose we want to measure how a particular type of optogenetic stimulus affected the lateral speed of the flies. More precisely, the user can specify the first .5 s of the 3rd, 4th, and 5th stimulus period over which to average the flies’ lateral speed and normalized this by the flies’ behavior in the 0.5 s before each of those stimulus periods:

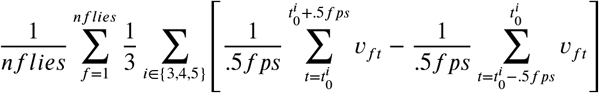

where 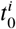 is the start frame of stimulus *i* and 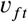 is the lateral speed for fly *f* at frame *t*. This allows the user to define the behavioral quantification that is most relevant to their experimental questions and design. These definitions are stored in the configuration files for the *feature statistics* and *summary plots* stages of the pipeline, and can easily be reused and modified as experimental conditions are modified.

For large high-throughput screens, data quality control is essential. We have engineered quality control checks into both our data acquisition software and our analysis pipeline. During the data acquisition, the experimenter can flag any potential issues with the experiment in its digital metadata. The pipeline checks that all data collection files exist at the start and that all expected output files have been generated by the end (Incoming quality control checks and Completed quality control checks).

### Multifly Pose Tracker: Dataset and Performance

The performance of supervised machine-learning algorithms is dependent on the quality of the training dataset, and modern deep learning algorithms perform best with large datasets. Here, we present *multifly*, a large pose estimation dataset for fruit flies enriched in complex social and motor behaviors, consisting of 9701 examples labeled with 21 keypoints on the head, thorax, abdomen, legs, and wings (***Figure 6, Table 1***, multifly dataset). This dataset is divided into a training set of 7208 flies and a test set of 2493 flies for evaluation. Annotators with fly domain knowledge were trained to reproduce reference labels to ensure consistency, accuracy and speed while labeling. They labeled flies performing a range of behaviors involving limb motion from walking and grooming to complex social interactions including male and female aggression and courtship. The annotators using temporal context and domain knowledge to determine the position of all keypoints on every example, including those that were self-occluded or occluded by social interactions, and labeled every fly within a given social interaction. We enriched the dataset with neural activation that increased the frequency and intensity of these social interactions. The annotators reviewed the tracker’s predictions, found errors, labeled by correcting the prediction, and added these high-value examples to the training dataset. We used our interactive Animal Part Tracker (APT) software to label the flies (***Kabra et al., 2022***). The APT user-interface was engineered to facilitate labeling: quick navigation between animals and across movies, multiple labeling modes, and iterative training and labeling. We used a two-stage, top-down method to train a multi-animal keypoint tracker where (1) targets are detected and (2) the keypoints are predicted for each target. Here, the body position and orientation trajectories from the FlyTracker were used as the first stage (***Eyjolfsdottir, 2017***). As shown in ***Figure 4***, a cropped and oriented image of each fly is passed the second stage where the x,y positions of each keypoint are predicted. We trained our multifly pose tracker with APT using the our GRONe (Grid Regression Offset NEtwork) algorithm. There is other pose estimation software including DeepLabCut and SLEAP commonly used by biologists (***Mathis et al., 2018***; ***Pereira et al., 2022***). In order to precisely quantify the performance of our tracker (***Figure 7***), we used our test set to compute the error between the tracker’s predictions and human’s labels. The test set was specifically designed to test both how well the tracker generalized to novel data and the effect of the flies’ behavior on keypoint prediction accuracy.

**Table 1.** Keypoint definitions.

| Num. | Name | Anatomical Description | Type |
| --- | --- | --- | --- |
| 1 | Anterior head | Midline anterior head between antennae | head |
| 2 | Right eye | Right posterior eye edge | head |
| 3 | Left eye | Left posterior eye edge | head |
| 4 | Left thorax | Left anterior-lateral prescutum | body |
| 5 | Right thorax | Right anterior-lateral prescutum | body |
| 6 | Posterior notum | Midline posterior scutellum | body |
| 7 | Posterior abdomen | Midline posterior abdomen | body |
| 8 | Right femur | Visible end of right mid-leg femur | femur |
| 9 | Right femur-tibia | Right mid-leg femur-tibia joint | femur |
| 10 | Left femur | Visible end of left mid-leg femur | femur |
| 11 | Left femur-tibia | Left mid-leg femur-tibia joint | femur |
| 12 | Right front-leg tarsus | Distal tarsal tip of right front-leg | tarsi |
| 13 | Right mid-leg tarsus | Distal tarsal tip of right mid-leg | tarsi |
| 14 | Right hind-leg tarsus | Distal tarsal tip of right hind-leg | tarsi |
| 15 | Left hind-leg tarsus | Distal tarsal tip of left hind-leg | tarsi |
| 16 | Left mid-leg tarsus | Distal tarsal tip of left mid-leg | tarsi |
| 17 | Left front-leg tarsus | Distal tarsal tip of left front-leg | tarsi |
| 18 | Right wing tip | Right wing edge where medial vein intersects | wing |
| 19 | Right wing | Right wing edge where radial vein intersects | wing |
| 20 | Left wing tip | Left wing edge where medial vein intersects | wing |
| 21 | Left wing | Left wing edge where radial vein intersects | wing |

**Figure 6.**
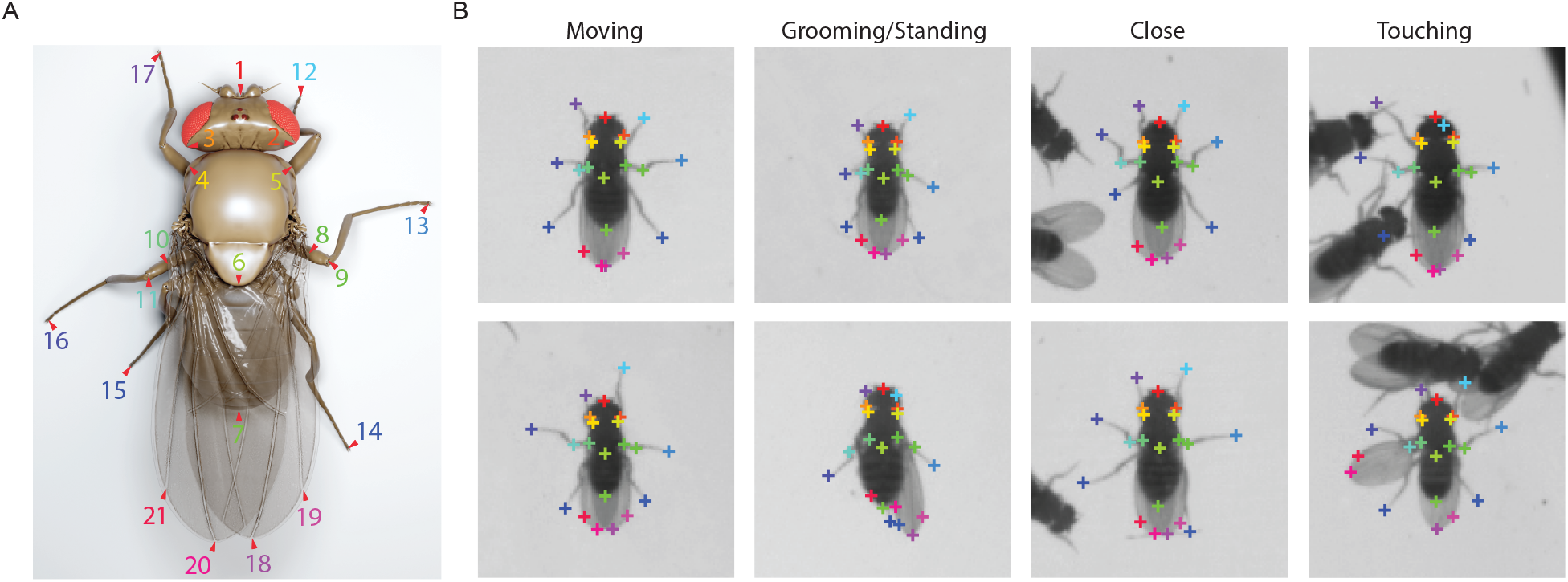
Keypoint definitions and example labels. (A) Illustration of the 21 keypoint locations on a rendered fly model. (B) Two randomly selected examples of manually labelled key points from each label category of moving, grooming/standing, close, and touching. Moving flies are translating and more than 3mm from any other fly, grooming/standing flies are not translating and more than 3mm from any other fly, close flies are withing 1.2-2mm of another fly but are not making physical contact, and touching flies make physical contact with another fly.

**Figure 7.**
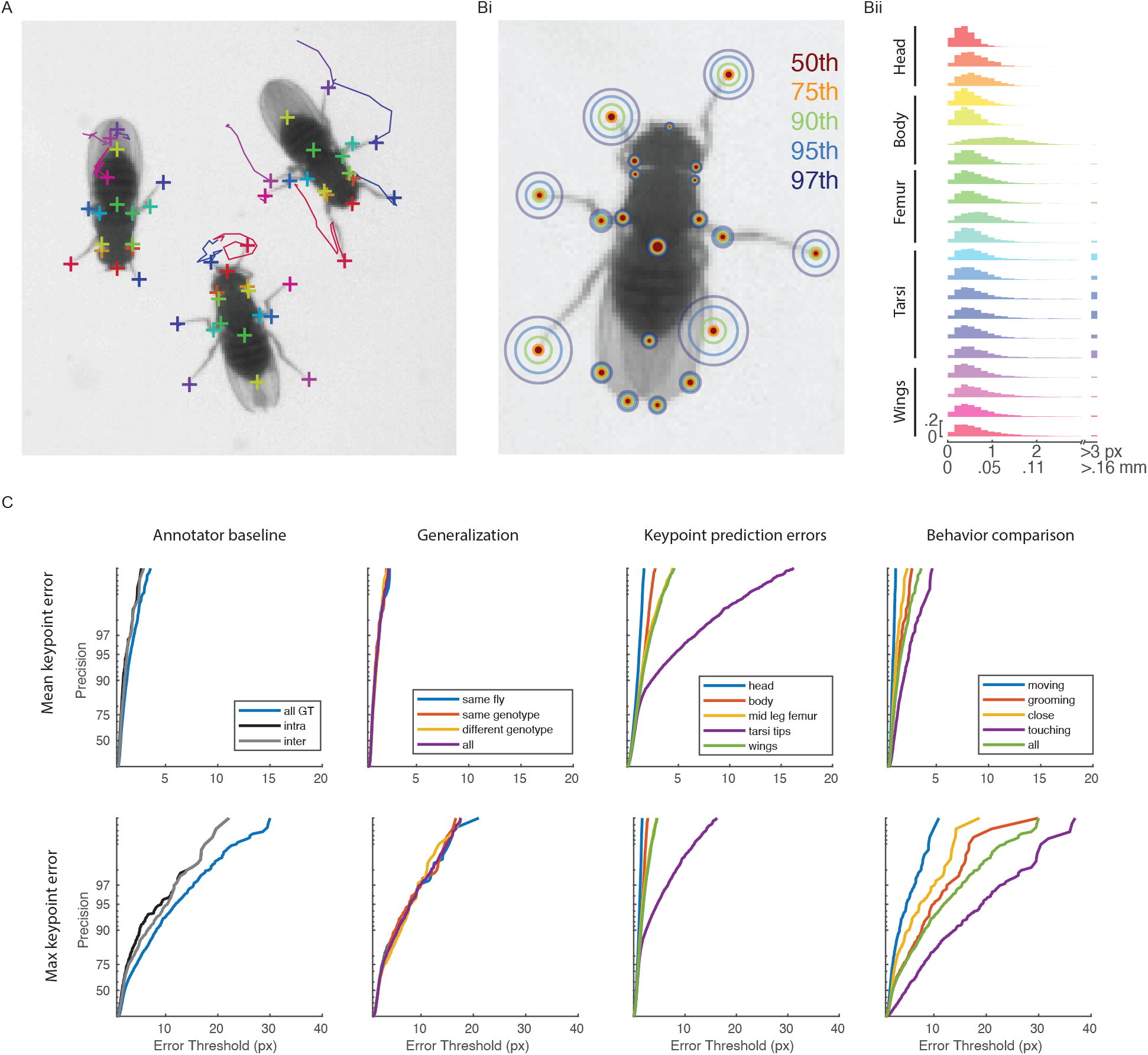
Keypoint tracking accuracy. (A) Example keypoint predictions. (Bi) Prediction errors for the test dataset (n = 2493) plotted as (i) circular histograms at the 50th, 75th, 90th, 95th, and 97th percentiles, and (ii) histograms for each keypoint grouped by type. (C) The precision of mean (top row) or maximum (bottom row) error of each example’s 21 keypoints at increasing error thresholds. The precision is shown for a different sets of data in each column: *Annotator baseline* is a comparison of the tracker’s predictions (n = 2493) to manual relabeling by the same labeler (intra-annotator, n = 597) and different labeler (inter-annotator, n = 597). *Generalization* compares how well the tracker works on increasingly novel subsets of test dataset: (1) examples sampled from the same flies as in the training dataset (same fly, n = 600), (2) examples from the same genotypes used in the training dataset but from different videos (same genotype, n = 600), (3) examples from different genotypes than the training dataset (different genotype, n = 600), and the entire test dataset (all, n = 2493). *Keypoint prediction errors* compares keypoints grouped by body part across the entire test dataset, and *Behavior comparison* show the effect of the flies’ behavior on tracking performance for flies that were (1) moving and not close to other flies (moving, n = 600), (2) had low velocity, usually grooming or standing still, and not close to other flies (grooming, n = 600), (3) within a body length of another fly but not in physical contact (close, n = 600), (4) in physical contact with another fly (touching, n = 693), and (5) the entire test dataset (all, n = 2493).

Our keypoint tracker is accurate: for our test set enriched for difficult poses, the mean error over all keypoints was .04 mm (1.5% of body length (BL)), with 95% of keypoint predictions within .11 mm (4% of BL) and 97% within .15 mm (5.6% of BL) (***Figure 7***). The tracker performed best on the keypoints of the head, wings, and thorax (excluding the notum) which had sub-pixel mean error (<.05 mm), and 95% within .08 mm. The leg tips, while more challenging, had a mean error of .06 mm and 95% within .24 mm. The tracker also generalized extremely well; we did not find any performance differences for prediction on novel flies or genotypes. Next we quantified the effect of body part and behavior on pose predictions. Fly legs have 5 actuated joints and 8 degrees of freedom without including the tarsal segments (***Lobato-Rios et al. (2022***); ***Vaxenburg et al. (2024***); I. Siwanowicz, personal communication) enabling a large range of limb positions. As expected from this variability in pose, leg-tip position predictions had lower precision than head, thorax, abdomen, and wing keypoints. Prediction precision depended on the behavior the fly was performing. Poses during social interactions were more difficult to accurately predict than during grooming, which were more difficult than while walking. The per-frame pose predictions have not yet reached the level of human annotators: the most accurately reproduced labels were produced by one human annotator repeatedly labeling the same examples followed by a different human relabeling. However, even two humans can have >20 pixel differences in their estimations of a keypoint, highlighting the challenge of pose detection especially during complex social interactions.

### Multifly Keypoint Tracking-based Behavior Analysis

High-quality keypoint trajectories are useful for fine-grained quantification of animal behavior. Here, we demonstrate using landmark positions as both the input to behavioral classifiers and to quantify locomotor behavior. We extended our behavioral classification software, JAABA, to use keypoint trajectories as input (***Kabra et al., 2022***). Based on these, JAABA computes a suite of engineered features for keypoints that includes metrics based on a single keypoint (either in the global or animal-centered reference frame), and user-defined pairs and triads. Keypoint pairs and triads were from a single animal as well as pairs of nearby animals, to detect social behaviors. We created accurate behavior classifiers for an individual behavior, *anterior grooming*, and a social behavior, *front-leg social touch* (***Figure 8***A,B). We defined anterior grooming to include cleaning the head parts with the forelimbs and the associated leg rubbing (***Seeds et al., 2014***). We defined front-leg social touch as a fly touching the body of another fly with one or both of its front legs. Neither of these behaviors could be accurately classified without accurate leg tracking (***Kabra et al., 2013***). For anterior grooming, the keypoint-based features were sufficient to generate a high quality classifier, 97.3% balanced accuracy, whereas for front-leg social touch, both body- and keypoint-based features were required to create an accurate classifier, 95.5% balanced accuracy.

**Figure 8.**
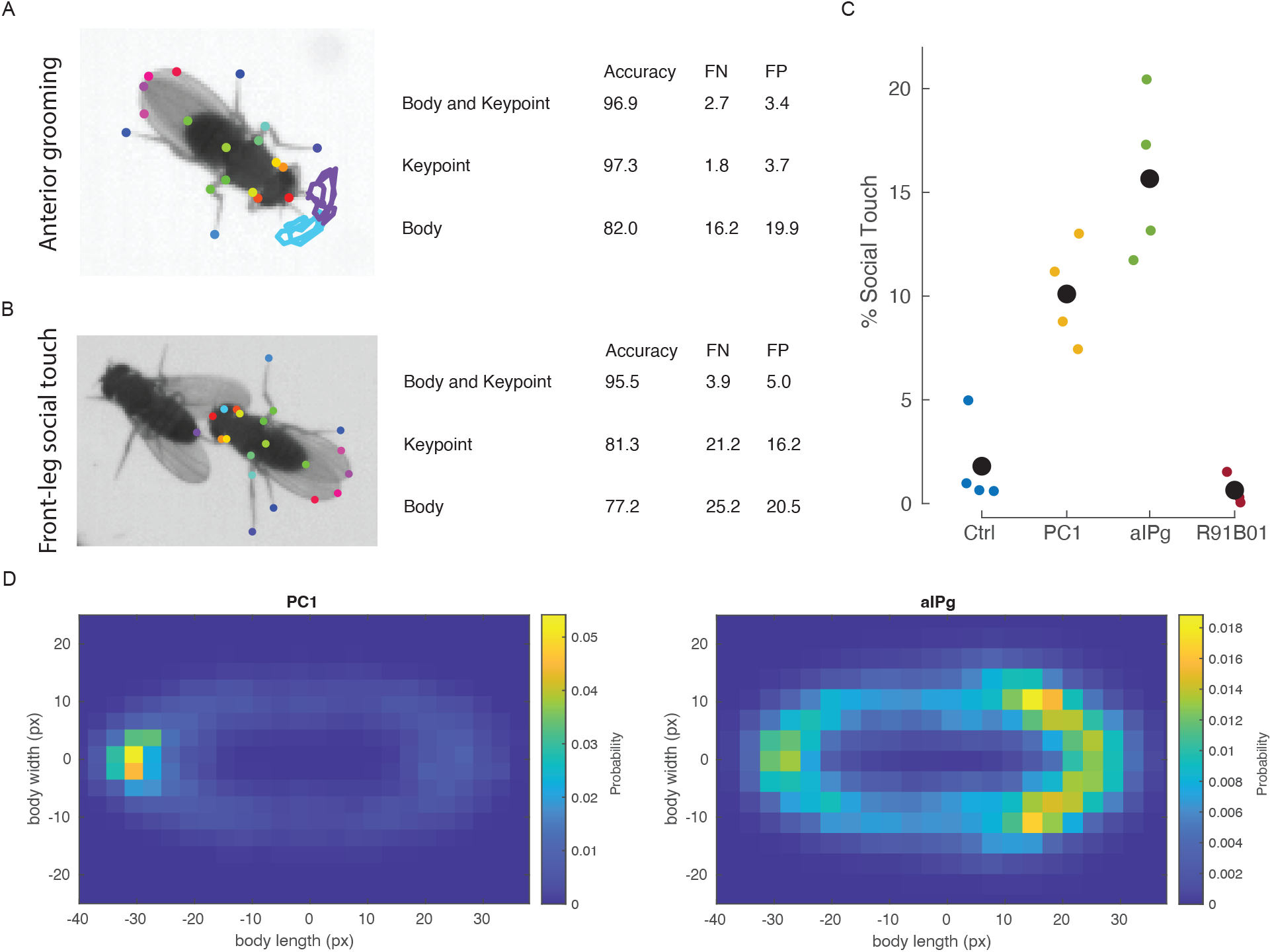
Behavior classification from keypoint trajectories. (A) An example of *anterior grooming* with keypoint tracking overlaid, and (B) An example of *front-leg social touch* with keypoint tracking for the classified fly overlaid. For each behavior, we trained classifiers and report the behavioral accuracy, the false negative rate and false positive rate computed using 7-fold cross validation. (C) The per video (colored dots) and total (black dot) averages of the percent of time the flies performed social touch during TrpA activation of empty-GAL4 control (5m,5f), pC1 line R71G01 (5m,5f), aIPg line SS36564 (10f), and R91B01 (5m,5f). For R71G01, only the behavior of the males was included. N = 4 movies, fly n = 40, 20, 40, 40 respectively. (D) For all social touches, we measured the location of the touching leg in the target fly’s coordinate system and computed the probably density map. The target flies are centered at 0,0 with the head pointing to the right.

We applied the social touch classifier to examine whether flies preferentially investigate different parts of the other fly in different social contexts. Activation of neurons involved in courtship and aggression causes flies to perform more social touches than control flies or flies with activation of avoidance neurons (***Figure 8***C). During front-leg social touch frames, we measured the location of the closest front leg tip in the target fly’s coordinate system. We observed differential targeting patterns for male pC1 activated flies and female aIPg flies (***Figure 8***D). The male flies almost exclusively touched close to the abdomen tip of the other flies. Aggressive females, by contrast, targeted their touches predominately to anterior region of their conspecifics but also targeted touches more generally around the entire body. The females are targeting touches to the posterior abdomen more than might be expected from their reported body orientation during aggressive behaviors ***Schretter et al. (2020***).

We developed a novel stage for the analysis pipeline, *locomotion metrics*, to quantify the kinematics of walking behavior in flies from keypoint trajectories. This stage can be used to quantify subtle changes in leg coordination. In ***Figure 9***, we show these metrics for control, and validate that these metrics match previously reported patterns (***DeAngelis et al., 2019***; ***Wosnitza et al., 2013***; ***Mendes et al., 2013***; ***Pratt et al., 2024***). We first computed whether a limb was in the swing or stance phase of locomotion using a simple state-dependent double threshold rule on leg tip velocity (***Figure 9***A). Taken together, the swing and stance state of the six limbs define the coordination pattern the fly is using to walk. Two coordination patterns have often been described, tripod (alternation of three legs swinging together) and tetrapod (a sequence of two legs swinging together). For control flies, we performed a simple per-frame classification of these patterns (***Mendes et al., 2013***) during walking. As expected, the flies in the bubble more often use the tripod pattern as they walked faster (***Figure 9***B). The flies modulated the duration of the stance phase of locomotion to facilitate faster movement but not the swing phase (***Mendes et al., 2013***; ***DeAngelis et al., 2019***; ***Pratt et al., 2024***). In ***Figure 9***C, we show the stance phase decreased as speed increased in each pair of limbs. Additionally, green LED light does not effect the behavior of these control flies – an important baseline for optogenetic experiments.

**Figure 9.**
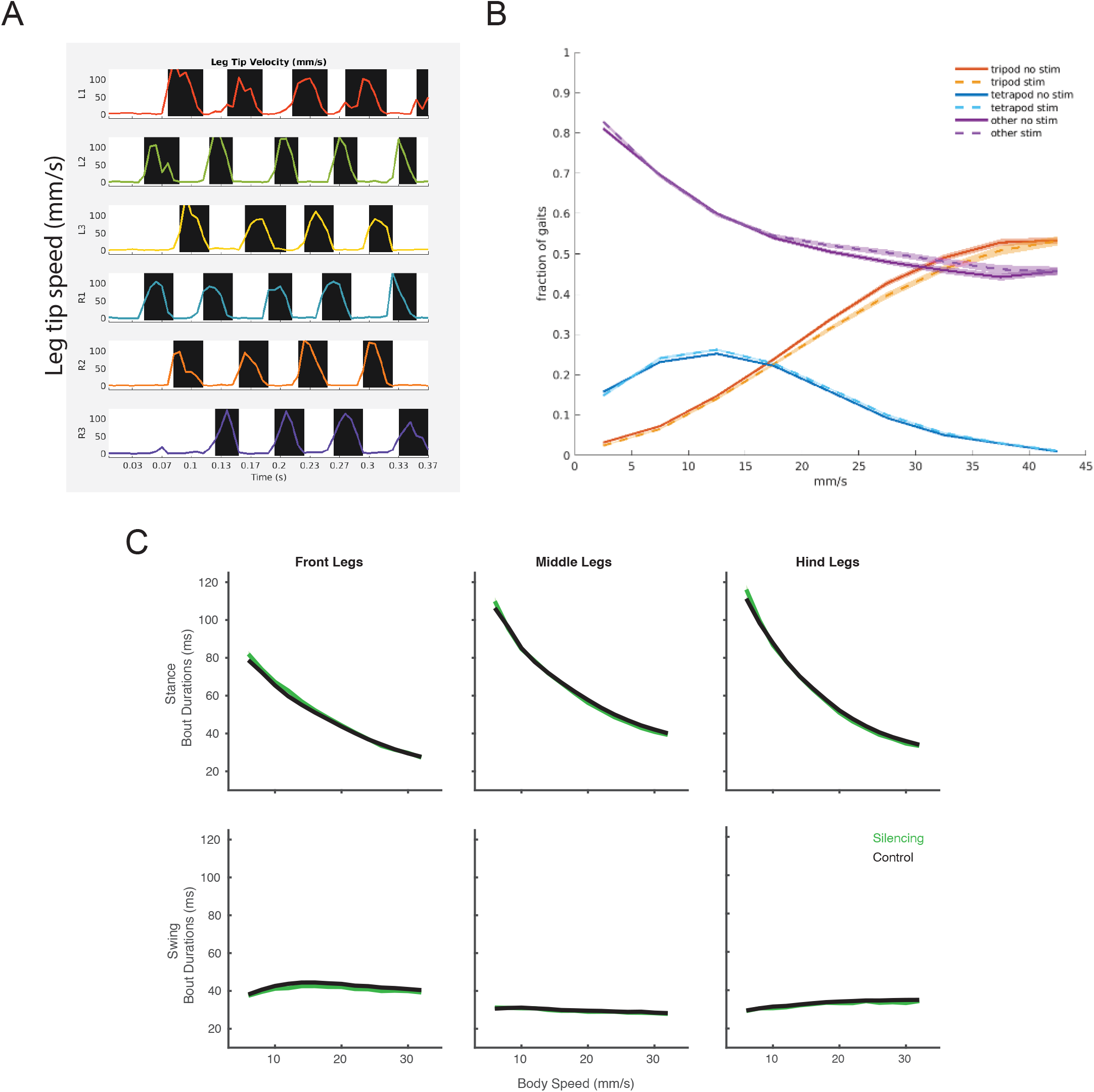
Locomotion kinematics. (A) Swing (black) and stance (white) overlaid on leg tip velocity for the L1 (left front), L2 (left middle), L3 (left back), R1 (right front), R2 (right middle) and R3 (right back) legs. (B) The fraction of locomotor patterns (tripod (orange), tetrapod (blue), and all other patterns (purple) for each bin of body speed during walking with no optogenetic stimuli (solid line) and green light (dashed). (C) The duration of each swing and stance bout during walking is plotted against the mean body speed for that bout. The data for the front, middle, and hind leg pairs are plotted from 8 split-GAL4 control videos when there was no optogenetic stimuli (black line) and green light stimuli (green line).

## Discussion

Here, present an end-to-end system for behavioral experimentation, from data collection through fine-grained behavioral analysis, for groups of freely walking and interacting fruit flies. The data acquisition system standardizes high quality video collection during optogenetic manipulation of neural activity. Our Fly Disco Analysis pipeline automates the machine vision- and learning-based analysis of the videos, including body and pose tracking, and produces a summary of the behavioral consequences of neuronal manipulations. In conjunction with brain-wide wiring diagrams and cell-type specific genetic reagents, this system can help scientists discover the functional role of neurons and circuits.

The system is composed of independently useful parts: the behavioral chamber design, the multicolor, programmable LED array, the analysis pipeline including keypoint tracking based on the multifly pose dataset, optogenetic-specific summary statistics and data visualization, and behavior classification based on JAABA extended to input keypoint trajectories. From the detailed documentation and fabrication instructions, our bubble chamber has been replicated by colleagues both within and outside of Janelia. Versions of the optogenetic LED array and backlight have been used in multiple behavioral studies (***Aso et al., 2014***; ***Dag et al., 2019***; ***Schretter et al., 2020***; ***Sawtelle et al., 2024***; ***Schretter et al., 2024***). Together with our collaborators, we have adapted the pipeline to analyze videos collected from other behavioral assays (***Schretter et al., 2020***), including a standard configuration of a single camera recording an array of small chambers for the study the behavior of single or paired flies. While the specific experimental design of our end-to-end system may not capture all aspects of fly behavior, each of these tools serve as an important resource for a vast array of behavioral experimental designs.

Our multifly dataset for pose estimation is a resource for both the neuroscience and the machine learning communities. Neuroscientists can use this dataset as a base and train trackers for substantially different setups through transfer learning (***Hoffman et al., 2018***; ***Doersch and Zisserman, 2019***) or fine tuning (***Ye et al., 2024***). Our pre-trained tracker was sufficiently general to work well in a similar setup, presumably aided by data augmentation built in APT. Pose estimation continues to be an active area of research in the machine learning community, particularly for difficult examples including occlusions and intense social iterations. Our multifly dataset is purposefully enriched for these challenging examples, and can enable new algorithmic development. Our annotators used the surrounding video to determine of location of limbs during complex social interactions or behaviors that occluded the limb. However, current pose recognition algorithms primarily operate on single frames. Incorporating temporal context may improve performance.

## Methods and Materials

The detailed parts list, 3D design files, and assembly instruction for the Fly Disco hardware are publicly available in our GitHub repository, https://github.com/arobie/FlyDiscoHardware. Links to all additional GitHub repositories subsequently referenced are also provided in the README file of the FlyDiscoHardware repository.

### Fly Disco Hardware

#### Fly Bubble behavioral chamber

The assembled Fly Bubble cartridge created the chamber where flies were recorded (***Figure 1***). The cartridge sandwiched the clear acrylic *window* between the 3D-printed *frame* and *scratch guard*, and was permanently screwed together along three sides. This created a slot for the *bubble* (press-fit into the *insert*) to be slid into the cartridge with the window forming the floor of the recording chamber and the bubble the ceiling. Importantly, the 3D-printed frame contained elastic elements that hold the bubble and insert firmly in place ensuring consistent positioning of the bubble in each video. This design allowed the bubble to easily slide in and out of the cartridge for loading and unloading flies and for the chamber to be quickly disassembled for cleaning between trials. Additionally, the cartridge included an asymmetry in its footprint matched in the mounting frame to prevent improper placement under the camera.

The bubble, the ceiling of the recording chamber, is a thin plastic sheet vacuum-formed to create a flattened shallow dome 3.5 mm tall in the center and 50 mm in diameter with rounded sloping sides (adapted from ***Berman et al. (2014***)). The interior of the bubble is coated with Sigmacote^®^ (Sigma-Aldrich), forming a microscopic silicone layer. This creates a slippery surface that reduces flies walking on the ceiling. The bubble is press fit to the bottom of the 3D-printed black insert with the bubble protruding through the large central hole. The black edges of the insert create high-contrast arena edges used by the pipeline to automatically detect the arena position.

The window, the floor of the recording chamber, was clear acrylic. The raised scratch guards of the cartridge help prevent the bottom surface from being scuffed. Silicon tape wrapped around the window facilitated the bubble sliding in and out of the cartridge and reduces scuffing of the window from above. Outside of the camera field of view, a countersunk hole in the window provided access to the recording chamber for loading flies. To load the flies, the assembled cartridge was placed upside-down in the loading frame. The bubble was then slid under the loading hole using the angled tab on the insert as a handle. The flies were transferred to the chamber with either our custom loader or via mouth pipette through the loading hole, and finally the bubble was slid firmly into place, enclosing the flies in the chamber. The flies were removed from the chamber using CO_2_ blown in through the loading hole to anesthetize the flies, and then they were suctioned out with a high velocity handheld vacuum or a vacuum line with ethanol trap. The window was cleaned with 70% ethanol between each trial. Bubbles were washed and Sigmacote^®^ (Sigma-Aldrich) reapplied weekly or more frequently if the flies were walking on the ceiling. The bubbles can be lightly wiped to remove dust, but mechanical abrasion damages the silicone layer. Every effort was made to remove dust and prevent scratches to the bubble and window. These degrade the quality of the images and the subsequent machine-vision analysis. To maximize data collection throughput, we built two cartridge assemblies per rig. This allowed the experimenter to unload, clean, and dry one chamber while the next video was being collected with the other chamber.

#### LED array

The LED array was designed to be both an IR backlight for video collection and a programmable optogenetic stimulus generator. The IR channel output constant illumination at a set intensity while the color channels were programmed with either constant or pulsed stimulation patterns. For further details of the LED controller, see LED controller. To remove excess heat and prevent heat damage to the LEDs, the LED array was mounted on a water-cooled breadboard. The coolant was circulated through a CPU-cooling radiator. A 1mm thick white acrylic sheet mounted above the backlight served as a diffuser. The board has had a number of design iterations for optogenetic stimulation including: single color boards, tricolor LEDs, and the current design which has three independent color channels. This allowed the use of red LEDs with longer wavelengths for example, or doubling the number of green LEDs to improve light output levels.

#### Fly Disco rig hardware

To create a compact footprint, the entire rig was built on a tall, narrow cart constructed of extruded aluminum which served as a superstructure for mounting hardware components but also held the computer, power supplies, and radiator. The flies in the bubble chamber were recorded at 1024 × 1024 pixels and 150 fps using a USB3 camera, FL3-U3-13Y3M (Point Grey). The camera was mounted to allow 3D adjustment of its position using a X-Y linear stage for fine adjustments left-right and linear rail for adjustments in the Z dimension. The camera was fitted with 25mm 1” format lens (Edmund Optics #63-246) and a 760nm IR pass filter (Hoya). The pass filter allowed wavelength separation between the imaging and stimulation illumination such that the video was unaffected by the optogenetic stimuli facilitating high quality background subtraction-based animal tracking. We developed a method for visualizing the time course of optogenetic stimuli in the video that did not negatively affect tracking. We added *indicator LEDs* to the system, these are IR LEDs placed outside the behavior chamber but still within the field of view of the camera. There was one IR indicator LED per color channel on the board. A small circuit board for the indicator LEDs was mounted to the bottom of the *baseplate* and aligned to the *mounting frame* for the bubble cartridge. This circuit board was connected to the LED controller via a spring-loaded contacts which allowed the baseplate to be removed without disconnecting wiring. The bubble cartridge insert has a small oval hole aligned to the indicator LEDs for visualization in the camera field of view. There were an additional pair of holes in the insert used for spatial registration through which the IR backlight was always visible.

To control the visual environment that the flies experienced, the behavior chamber was enclosed on all 4 sides with blackout panels. The front panel was removable for loading the flies and was attached with magnetic latches for easy of use. A second set of latches provided an *open* position used primarily during the fly loading procedure. This visual surround isolated the behavior chamber from visual distractions of experimenters in the room. The top of the cart was open to facilitate air exchange with the temperature and humidity controlled room. As flies use vision information for stabilizing locomotion (***Robie et al., 2010***) and during social interaction, we provided dim white light, 20-30 lux) to the flies via four voltage-regulated LED lamps diffused by a 1mm white acrylic sheet mounted above the behavioral chamber.

#### LED controller

The LED controller used an Arduino^®^ that received the high-level commands from data acquisition software (FBDC) via a USB-Serial command interface. The static intensity level of the IR LEDs was specified via FBDC and sent to the LED board controller during initialization of the FBDC GUI. The IR LEDs were controlled via pulse-width-modulation to minimize overheating, but this also caused the floor effect seen in their power-intensity curve (***Figure 3***B). For further details of FBDC see Data acquisition software.

For the color LED channels, the user created the optogenetic stimulus pattern in a simple Excel^®^ spreadsheet. The stimulus protocol file is organized by *steps* of LED activation patterns. Within each step the same pulsed pattern is repeated a given number of times at the same intensity. As illustrated in ***Figure 10***, the LED activation pattern of each is parameterized by pulse width, pulse period, the number of pulses per iteration, the length of the pause between iterations, and the number of iterations. Within a step, the pulse patterns for each color LED can be different, but any pattern that is longer than the step duration will be truncated. FBDC interpreted in the LED protocol file and passed the individual step commands in the controller. The controller used linear analog control of current to drive the timing and intensity of individual RGB LEDs.

**Figure 10.**
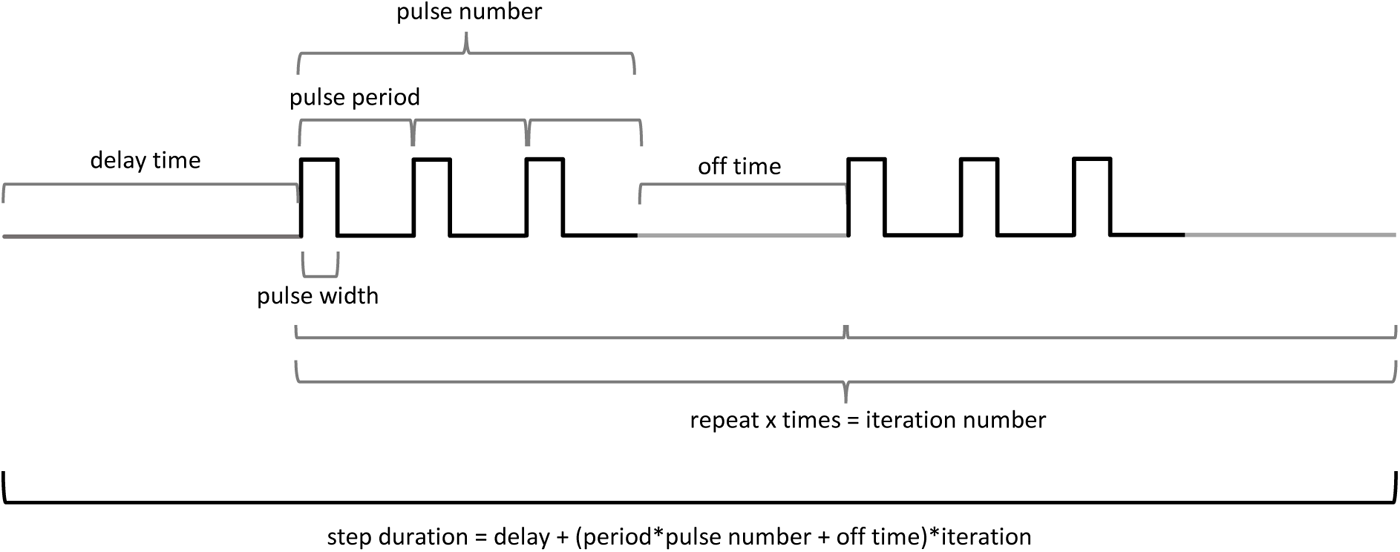
LED stimulation parameterization. Graphical illustration of the values used to parameterize the LED stimulation protocol for a single step of one color LED in the stimulus protocol file.

The LED controller also drove the indicator LEDs: the three IR LEDs in the camera field of view that display the activity of RGB LEDs respectively. These LEDs displayed a low pass version of the stimuli e.g. the envelop of the stimuli profile, to avoid sampling issues during short duty cycle pulsed stimuli. This allowed the classification of the stimuli status for each frame directly from the video and the recovery of the optogenetic stimulation pattern for analysis (Indicator LED detection).

The publicly available GitHub repository for the LED boards (https://github.com/janelia-experimental-technology/RGB-IR-LED-Boards) contains all the necessary components for manufacture and use: the specification and parts list for fabrication for multiple LED board designs, the common firmware code, and a description command interface. The firmware and command interface is common across all versions of the LED board. The current designs are 5”x5” and 7”x7” with 3 independent LEDs channels in addition to the IR channel. As shown in ***Figure 2***A, groups of IR and color LED channels evenly tile the board. Each of the four quadrants of the LED array can be independently controlled, and multiple boards can be daisy-chained together using a common controller.

### Data acquisition software

Our data acquisition software was designed to facilitate high throughput data collection while minimizing human error. This MATLAB software GUI controlled the video collection, the optogenetic stimuli, and the collection of metadata information about the experiment such as the driver line name and room temperature. We made a number of significant updates to the FlyBowlDataCapture software (***Robie et al., 2017***) that include the acquisition of movies from USB3 cameras using our BIAS software with either the FlyCapture or Spinnaker SDKs, and the integration of control for the LED boards. All the software is freely available on GitHub: FBDC at https://github.com/kristinbranson/FlyBowlDataCapture and BIAS at https://github.com/kristinbranson/BIASJAABA

The data acquisition GUI is initialized with a parameters file (FBDC parameter file) that specifies the details of how a given experiment will be run including the camera configuration file, the intensity of the IR backlight LEDs, the file location of the optogenetic stimulation protocol for the red, green, and blue LEDs, the ranges for metadata values, temperature and humidity recording frequency, recording time period for non-optogenetic experiments, and the computer locations for additional parameter files and a storage location for the experimental data (***Figure 3***).

After the user loads the FBDC parameter file, the FBDC GUI opens with pre-populated fields based on the parameter file’s ranges and the defaults from the last use of the GUI. These values should be adjusted to specifics of a given experiment and will be stored in the digital *metadata file* created for each video. The user presses the *initialize camera* button to launch an instance of BIAS which will open with the camera running in preview mode. The user then loads flies into the bubble, places the assembled cartridge into the rig, replaces the front panel, and presses the *Flies Loaded* button to capture the timestamp of this action. To start the trial, the user presses *Start Recording*. A delay, to allow flies to settle down from the loading transfer, before video acquisition can be specified in the FBDC parameter file. After this delay (if any) the GUI signals to BIAS to start recording the video and to the backlight controller to start running the LED stimuli protocol. For non-optogenetic experiments, the movie length is specified in the FBDC parameter file, and for optogenetic experiments the move length is calculated from the LED protocol. The BIAS window will continue to display a live preview of the video throughout the experiment, and the FBDC GUI will display the progress through the optogenetic stimuli, the real-time frame rate of the camera, and experimental status updates. Throughout the experiment, the user can update the metadata fields as needed, record technical or behavioral notes, and flag experiments. Additional fields allow the documentation of the number of damaged or dead flies in the bubble. Using the *Abort* button, the experiment can be terminated at any point. To facilitate flagging any quality control issues, after the video collection finishes the software displays the recorded video and a technical summary of the video collection and compression performance. Any metadata field can by updated until the experiment is ended by pushing the *Done* button. A new experiment can be quickly started using the same FBDC parameter file by selecting *new experiment* in the file menu and re-initializing the camera.

All experiments using the same experimental parameters such as LED stimulation pattern and movie length can use the same FBDC parameter file. The GUI will capture the experimental difference between flies in the metadata file through the *experiment* and *condition* fields. These drop-down fields are populated by condition parameter files in the directory specified by the *ConditionDirectory* parameter in the FBDC parameter file. The names of these files must follow the format *Experiment_Condition*.csv. If there is a single experimental variable per video such as driver line or starvation length, this can be represented by the condition field. In the case where the condition is the driver line, such as in an optogenetic screen, the condition field can be populated using a barcode scan or from a list of lines in the condition parameter file. In these cases the condition parameter file must follow the format *Experiment*_BARCODE.csv. If the experimental design is multi-factorial such as three starvation lengths for experimental and control fly lines, then a condition file for each pair of factors is required.

### Fly Disco Analysis Pipeline

A helper program, Transfero (https://github.com/JaneliaSciComp/transfero), runs nightly as a Linux `cron` job. Transfero communicates with the set of computers used for data collection, and scans each for new experimental data. The data is organized (by the data acquisition software) into *experiment folders*, each containing the data from a single experimental video. Any new experimental folders are copied to a central scale-out storage system (Vast Data) via the `scp` protocol unless the experiment was aborted. Aborted experiments are deleted from the data collection computer and not copied to the storage system. An MD5 checksum is computed on each data file at source and destination to ensure data integrity. After transfer, the Fly Disco Analysis pipeline is run on each experiment folder, each as a single LSF (IBM) job on a local HPC cluster.

The Fly Disco Analysis pipeline is implemented primarily in MATLAB (Mathworks). It is freely available on our GitHub page, https://github.com/kristinbranson/FlyDiscoAnalysis. Additional required tools such as FlyTracker, JAABA, and APT are included as submodules of the FlyDiscoAnalysis repository. The keypoint tracking (APT) stage requires access to a GPU to perform deep-learning based inference, and is implemented in Python.

The FDA pipeline consists of a number of individual *stages*, where each stage’s input is provided by the output of some previous stage(s) (***Figure 5***). Furthermore, some stages are optional, and their output is only required by subsequent stages if they are turned on (colored stages in ***Figure 5***). Because some stages are expensive to run, the FDA pipeline is designed to be ‘resumable’, and each stage is generally only run if that stage’s outputs are missing. Specifying which stages are run and the parameters for those stages is done in a settings directory that contains configuration files for the pipeline and stage-specific parameter files. Each stage can be be set to *on* (run if the output files for that stage do not exist, *off* (do not run), or *force* (always run, overwriting preexisting output files if needed). Each experiment folder contains a `metadata.xml` file that specifies the *screen type* of the experiment. The screen type is used to determine the settings folder used for that experiment. In ***Table 2***, we enumerate each stage of the pipeline and its input parameter files and output files.

**Table 2.** The stages of the pipeline with input and output files. (1) Each stage of the pipeline and the name of its source code, (2) input parameter files, (3) when the parameter files likely need to be edited either during the initial set-up of a new rig or for new experimental design including which parameters should be updated, and (4) the output files of each stage.

| pipeline stage and code | input parameters files | edit for bubble experiment | output files |
| --- | --- | --- | --- |
| **incoming checks**<br>FlyDiscoAutomaticChecksIncoming.m | automatic_checks_incoming_params.txt | new exp - expected movie length | automatic_checks_incoming_results.txt<br>automatic_checks_incoming_info.mat |
| **body tracking (FlyTracker)**<br>FlyDiscoFlyTrackerWrapper.m | parent_calibration_file.mat<br>flytracker-options.txt | initial set-up | movie-track.mat<br>flytracker-calibration.mat<br>movie-bg.mat<br>movie-feat.mat<br>movie-params.mat<br>movie_JAABA/trx.mat<br>movie_JAABA/movie.ufmf (softlink) |
| **registration**<br>FlyDiscoRegisterTrx.m | registration_params.txt<br>Registration_mark_template.png | initial set-up | registered_trx.mat<br>registrationdata.mat<br>registrationdata.txt<br>registrationimage.png |
| **indicator LED detection**<br>FlyDiscoDetectIndicatorLedOnOff.m | indicator_params.txt<br>LED_indicator_ON_template.png | new exp - optogenetic or not<br>initial set-up | indicatordata.mat<br>ledregistrationimage.png<br>indicator_log.txt |
| **sex classification**<br>FlyDiscoClassifySex.m | sexclassifier.txt<br>sexclassifier_params.txt<br>metadata.gender = b,m,f,x | initial set-up<br>new exp - metadata.gender | sexclassifier_diagnostics.txt<br>registered_trx.mat (modified)<br>sexclassifier.mat (sometimes) |
| **keypoint tracking (APT)**<br>FlyDiscoAPTTrack.m | apt_params.txt | initial set-up | apt.trk<br>apt_bsub_log.txt<br>apt_error.txt<br>apt_log.txt |
| **feature computation**<br>FlyDiscoComputePerFrameFeatures.m | perframe_params.txt<br>landmark_params.txt<br>perframefns.txt |  | perframe/*.mat<br>perframefeatures_info.mat<br>wingtracking_results.mat |
| **behavior classification (JAABA)**<br>FlyDiscoJAABADetect.m | JAABA_classifier_params.txt | initial set-up | scores_*.mat |
| **feature statistics computation**<br>FlyDiscoComputePerFrameStats.m | statflyconditions.txt<br>statframeconditions.txt<br>stats_params.txt<br>stats_perframefeatures.txt<br>hist_perframebins.mat<br>hist_params.txt<br>hist_perframefeatures.txt | new exp -all | stats_perframe.txt<br>stats_perframe.mat<br>hist_perframe.txt<br>hist_perframe.mat |
| **summary plots**<br>FlyDiscoPlotPerFrameStats.m | stats_plotparams.txt<br>hist_plotparams.txt<br>conditions_plotparams.txt<br>stimulus_plotparams.txt<br>stimulus_videoparams.txt | new exp - all | analysis_plots/hist_*.png<br>stats.html |
| **body tracking snippet movie**<br>FlyDiscoMakeCtraxResultsMovie.m | ctraxresultsmovie_params.txt or<br>ctraxresultsmovie_params_*.txt | new exp - snippet definitions | flytracker_results_movie.mp4 |
| **keypoint tracking snippet movie**<br>FlyDiscoMakeAPTResultsMovie.m | ctraxresultsmovie_params.txt or<br>ctraxresultsmovie_params_*.txt |  | apt_results_movie.mp4 |
| **completion checks**<br>FlyDiscoAutomaticChecksComplete.m | automatic_checks_complete_params.txt | new exp - *nflies, required_files | automatic_checks_complete_results.txt<br>automatic_checks_complete_info.mat |

The individual FDA pipeline stages are further described below.

#### Incoming quality control checks

To identify experiments with technical or user errors and prevent them from being further processed, we perform automated checks at the beginning of the pipeline. Experiments will be excluded from further analysis (“failed”) if (1) the `metadata.xml` file or movie file is missing, (2) the video file is too shorter than expected, (3) the fly loading time exceeds a specified length, (4) the experimenter manually indicated an error with data collection by aborting the experiment, setting a redo flag or setting the number of dead or damaged flies >0 (***Robie et al., 2017***). For optogenetic experiments, an experiment will also be failed if the stimulus protocol file is missing. Any of these errors result in an `automate_pf` value being set to ‘F’ and an `automated_pf_category` being set to the type of error in the *incoming checks* output files (see ***Table 2***), and the pipeline being terminated. If no errors are detected, the `automate_pf` value is set to ‘U’ (for ‘unknown’) and the pipeline continues.

#### Body tracking: FlyTracker

We track the flies’ bodies and wings with a modified version of the Caltech FlyTracker (***Eyjolfsdottir*** (***2017***), https://github.com/kristinbranson/FlyTracker). FlyTracker performs background-subtraction-based fly detection on each frame. The fly identities are constructed by linking detections across frames. The pixels near flies are then classified as body, wings, legs, or background. An oriented ellipse is fit to the body, line segments to the wings, and points to the leg tips. The pose is used to resolve potential identity swaps in the tracking. Important modifications we made to FlyTracker include: (1) adding a function to enable tracking without the GUI, (2) streamlining of the input parameters to simplify their specification in batch usage, and (3) improved head-tail disambiguation. Each analysis-protocol folder contains a *parent calibration* containing such parameters and the arena radii (in mm), the number of arenas in each video, and the shape of the arenas (circular or square). It also contains kernels used for segmentation and appropriate thresholds for distinguishing foreground from background. FlyTracker provides a tool for tuning these parameters based on an example video. Additionally, FlyTracker allows the user to set the number of CPU cores used for tracking, and the number of chunks the video is divided into for analysis. For consistency, we always used two cores and four chunks. (These parameters are specified in the `flytracker-options.txt` parameter file.) The FlyTracker stage outputs tracking data for the bodies and wings of the flies in each frame. This information is output in a JAABA-compatible format to permit subsequent analysis.

#### Registration: Spatial and temporal registration of tracking data

In the registration stage, the trajectory data from FlyTracker is spatially transformed: it is translated such that the coordinate system origin is the center of the arena it belongs to, and scaled to convert from pixels to millimeters. The boundaries of the arenas are determined either by 1) using the outlines provided by FlyTracker, or 2) using a Canny edge detector followed by a Hough transform to identify the circular arenas in the image (***Eyjolfsdottir, 2017***; ***Robie et al., 2017***). The black bubble insert that forms the arena is designed to be high-contrast against its light background, which aids in arena detection. The boundaries of the arenas are defined by the arena center and diameter in pixels. The arena radius in pixels together with the arena radius in mm (supplied by the user in parent calibration file) are used to define the pixels-to-mm scale factor. This spatial registration eliminates small differences in camera position across rigs and enables conversion of position data into real-world units of millimeters for subsequent computations such as velocity.

In addition to spatial registration, temporal registration is also done as part of the registration stage. Temporal registration of the trajectory data allows the user to select which sections of the video to analyze. This can be based on the time point the flies were loaded into the arena, if there is a variable time between video start and fly loading (***Robie et al., 2017***). Although this form of temporal registration is still supported in the Fly Disco pipeline, for the experiments reported here we added a temporal truncation feature, which allows the user to remove a defined number of frames from the end of the movie. This allows us to remove a small number of frames that were captured after the IR backlight LEDs were turned off.

One final computation is done during the registration stage. The trajectories output from FlyTracker sometimes contain IEEE-754 NaN values for some frames of a trajectory. (This can happen e.g. when two flies are close together, and their individual body pixels become difficult to discern.) In the registration stage, we remove these NaNs either by interpolating tracking data across short gaps (the maximum gap length is specified by the user) or splitting trajectories into multiple IDs when the gaps are larger.

#### Indicator LED detection

The indicator LED provides a reference timeline of the optogenetic stimuli in the video, eliminating the necessity of synchronizing digital inputs. Extracting the timing signal happens in two steps. First the location of the indicator LED in the video is detected and then the timing signal is extracted. The location of the indicator LED in the video image is automatically detected using template matching. The template is a tightly cropped image of the indicator LED in the *on* state. Based on the stimulus protocol, the algorithm creates a maximum intensity image, `ledMaxImage`, by taking the max of an evenly-spaced sequence of frames likely to contain many frames with the LED illuminated. If there are no pixels in this image as bright as those in the template, a new `ledMaxImage` is constructed, sampling from the entire movie without reference to the LED protocol. The location of the indicator is detected using normalized cross correlation of the template image and the `ledMaxImage`. A visualization of the detected location in the `ledMaxImage` is output to the experiment directory. The status of the indicator LED in each frame (illuminated or not) is computed using a threshold on the mean of the pixel intensity values in a box centered on the detected location of the indicator LED. This digital signal is processed to find the start and end frames of each LED activation bout, which are output in terms of both frame number and timestamps values. Finally, the number of detected bouts are compared in the LED protocol as a sanity check.

#### Sex classification

The sex of the flies is an important factor in the analysis of behavior. If mixed sex groups of flies are being recorded, automated sex classification based on the size of the flies assigns sex to each fly in each frame (***Robie et al., 2017***). The pipeline determines whether to run the sex classification algorithm based on the `metadata.xml` field *gender* which can have the values *b* (both males and females), *f* (females only), *m* (males only), or *x* (unsorted mixture). For *b*, the algorithm expects approximately equal proportions of males and females. In all cases other than *b*, the sex classification algorithm does not run, and all flies are directly assigned that value (m, f, or x) for sex. A diagnostic file is output that includes statistics on the number of total flies, males, and females detected in the video; the swaps in sex classification; and size of the flies. This diagnostic file is created regardless of whether the sex classification algorithm runs and provides the input for quality control checks on the number of flies that are run in the final stage of the pipeline.

#### Keypoint tracking: APT

Keypoint tracking is the automated detection of the position of an animal’s individual body parts. This enables behavior analysis based on the animal’s pose and limb movements. We use our supervised deep-learning software package, the Animal Part Tracker (APT) (***Kabra et al., 2022***), to detect 21 keypoints on the fly using a pretrained tracker (***Figure 6, Table 1***). These include three points on the head, three points on the thorax, the tip of the abdomen, the tips of each leg, the femur and femur-tibia joint of the middle legs, and two points on each wing. We employ a top-down tracking method, using the FlyTracker body tracking as the object detection step followed by single animal pose detection using the GRONe algorithm for each fly, in each frame, as illustrated in the part tracking inset of ***Figure 4***. Identity tracking has already been performed by FlyTracker body tracking, thus no individual linking step is required. The tracking accuracy and use of keypoint tracking is detailed in sections Multifly Pose Tracker: Dataset and Performance and Multifly Keypoint Tracking-based Behavior Analysis.

#### Feature computation

*Per-frame* features are a suite of metrics calculated from the body-tracking trajectory data that describe the animal’s behavior at a given frame. These continuous-valued metrics include locomotion features, such as the speed and rotation of the fly, and social features, such as the fly’s position and movement relative to other flies. The per-frame features are hand-engineered metrics that were developed as input for our automated behavior classifier JAABA (***Kabra et al., 2013***) and are described in detail in ***Robie et al. (2017***): Table S2. In this stage we compute them as potential input to higher-level JAABA classifiers, but also because they are useful descriptors of behavior and can be used to detect behavioral changes due to experimental perturbations.

#### Behavior classification: JAABA

We have updated our interactive machine learning software, JAABA, to train behavior classifiers based on keypoint trajectories (***Kabra et al. (2013***), https://github.com/kristinbranson/APT/wiki/Using-APT-tracking-data-in-JAABA). The per-frame features for keypoint data include locomotion, appearance, and social features for single, pairs, or triads of keypoints. The keypoint features can be used in combination with body-tracking-derived features or as the only input features to JAABA (useful in the case of keypoint tracking without body tracking). This stage of the pipeline applies pretrained behavior classifiers to produce a `score*.mat` file for each behavior. We have created an anterior grooming and a social touch classiers that required keypoint features for good performance. The specifics of our pretrained FlyBubble behavior classifiers in discussed further in Multifly Keypoint Tracking-based Behavior Analysis.

#### Feature statistics

We developed a controlled vocabulary to flexibly define metrics that describe the flies’ behavioral responses to experimental manipulations. This requires defining which frames and which flies to include in computations, and which metrics are being computed. This is done in the input parameter files of the feature statistics computation stage (***Table 2***). Briefly, the user can define frame periods to be analyzed based on the status of the stimuli (including partial periods relative to the beginning or end of stimulus periods that can be defined in seconds or percentages), behavior classification, or thresholds on per-frame features. The definitions take the form `<name>,<frame condition>,<state>`. For example, `stimon,equal_stimon,1` would include all frames when the optogenetic stimulus is active while `stimon,equal_stimon_1_2_3,1` would include only data from the first three stimulus periods. Similarly all frames where the front-leg social touch behavior is predicted can be defined by `touch,equal_labels_socialtouch,1`. To further refine the analysis period, these definitions can be combined: frames during optogenetic stimulation when the fly is performing the social-touch behavior. Next, the user can define which statistics will be computed using these frame periods and fly conditions. Examples include the female flies’ forward velocity, the flies’ turning rate in the stimulus period normalized by the turning rate in non-stimulus periods, or the fraction of time spent performing a behavior. This flexible system allows the user to tailor the analysis to (1) their specific experimental structure and (2) investigate behavioral changes relevant to their experimental questions.

#### Summary plots

To visualize the behavioral statistics computed in the previous stage, and to facilitate the exploration and discovery of behavioral phenotypes, this stage of the pipeline automatically creates a variety of summary plots. These plots include group and individual averages, time series, and stimulus-triggered averages of the behavioral metrics defined in the previous stage. We also create stimulus onset trajectory plots and images. These plots are programmatically generated based on user-defined configuration files. For ease of viewing the experimental results, a local web page is created to display the plots and movies.

#### Results movies for body tracking and parting

For visual inspection of tracking performance and behavioral responses, we create short MPEG-4 movies that display body (or keypoint) tracking. The movies display the full view of the chamber on the left and zoomed-in images of the flies on the right. These movies are composed of snippets from the full length video. The specifications of the snippets include whether they are centered on particular frames or onset of specific stimulus periods, and the number of frames before and after the centering point to include. We have found these movies to be invaluable for discovering unexpected behavioral responses, and also for reviewing tracking results when quality control metrics indicate a problem.

#### Completed quality control checks

The last stage of the pipeline is a final quality control check. To ensure the pipeline has properly generated the expected output files for each active stage of the pipeline, we verify the existence of the output files from every active pipeline stage. If any output files are missing, the experiment is marked as failed. We also record the category of failure and the names of the missing files. Here, a hierarchical system of categories ensures the earliest stage with missing files is reported as the cause of the failure, enabling quick troubleshooting of the pipeline. Additionally, the number of flies is checked to make sure it is within an acceptable range (specified by a parameter file). Similar ranges can be set for numbers of males and females. For optogenetic experiments, this stage also checks that the expected number of stimulus periods were detected from the indicator LEDs, to ensure that the stimulus protocol was properly generated and captured in the video. While it is likely that the wrong number of stimuli would have already caused an error in a preceding stage, this acts as final backstop against undetected errors in this important experimental parameter.

### multifly dataset

We created our labeled dataset, multifly, with trained annotators who had fly domain knowledge. Each annotator was trained to reproduce baseline example labels. Importantly, this training period helped correct systematic bias in positioning of labels between annotators and calibrated the annotators to the expected labeling accuracy and speed. The multifly dataset in the COCO data format is available at https://research.janelia.org/bransonlab/multifly.

#### Training set

We generated the training dataset in roughly 4 behavior-based batches (***Figure 6***B) using 9 independent videos of 9-10m flies from 8 different genotypes. The first batch of labels were from the easiest examples: flies were moving and not close to other flies, thus reducing occlusions. In the second batch, we labeled flies that were either grooming or standing still, and not close to another fly. During grooming, keypoints were often severely occluded. Every keypoint was labeled for every fly. However, when annotators had to infer the location of keypoint from temporal context or guess the location based on domain knowledge, they added an occluded flag to that keypoint label. In the third batch of labels, we included flies that were within a body length of another fly, but not in physical contact. Finally, in the fourth batch, we included flies in physical contact social interactions including male and female aggression and courtship. Every fly in physical contact within an interacting group was labeled. We excluded flies from the training dataset that were attempting to climb on the ceiling, walking on the ceiling, righting themselves from falling off the ceiling, jumping, or flying. To facilitate labeling of social interactions which would otherwise be quite rare in our experimental conditions (group-housed males and females), we artificially activated identified neurons with either TrpA or CsChrimson to induced social interactions.

#### Test set

We created a test dataset with the same annotators. We created the test dataset consisting of 2493 labeled examples to test how well the tracker generalized and how the behavior the fly was performing effected pose estimation. For the generalization, we annotated 600 examples in each of 3 categories: (1) flies used in the training dataset but from different parts of the movie, and flies from novel movies with either (2) same genotypes used in training dataset or (3) different genotypes. This allowed us to test the generalization performance on different degrees of novelty. Furthermore, in order to quantify the impact of the fly’s behavior on tracker performance, each generalization set contained 200 flies from each of the behavior categories: (1) moving, (2) standing/grooming, and (3) close. This resulted in 200 flies for each combination of generalization and behavioral category for a total of 1800 labeled flies. For the touching test dataset, we labeled 600 examples of flies involved in social interactions: 200 examples from lines with either induced aggression (male or female) or courtship phenotypes, and an additional 200 flies from lines without social phenotypes to include lower intensity social interactions. None of these movies were included in the training data. We also labeled every fly within the contact group for the focal fly, this resulted in 693 labeled examples.

#### Human baselines

An important baseline for supervised machine learning algorithms is how well human relabeling compares to predictions, i.e. does the algorithm reach human performance? We relabeled a subset of our test dataset to investigate human labeling accuracy. Either the original annotator relabeled (intra-labeler) or a different annotator relabeled (inter-labeler) the example. The same subset of the test data was relabeled in both cases: 597 flies spanning the behavioral and generalization categories with ∼150 labels per behavioral category (moving, grooming, close, and touching).

### Animals

Flies were sorted into groups of 5 males and 5 females or 10 females for SS36551, SS36564, and SS56987 under cold anesthesia at least 24 hours prior to behavioral experiments. The genotypes of flies used in the training, test, and APT behavioral experiments are listed in ***Table 3***.

**Table 3.** Genotypes used in the multifly dataset and behavioral experiments.

| Genotype | Source | Identifier |
| --- | --- | --- |
| *D. melanogaster*: 20XUAS-CsChrimson-mVenus (attP18) | Klapoetke et al., 2014<br>Aso et al., 2014 | CsChR |
| *D. melanogaster*: +; UAS-dTrpA1/CyO | Hoopfer et al., 2015 | TrpA |
| *D. melanogaster*: w1118;; 20XUAS-GtACR1-EYFP (attP2)/TM2 | Barry Dickson/Ryo Minegishi | GtACR |
| *D. melanogaster*: vGlut-GAL4 | Deng et al., 2019 | vGlut |
| *D. melanogaster*: w1118;; BPZpGAL4DBD (attP2) | Pfeiffer et al., 2010 | Empty GAL4 |
| *D. melanogaster*: w1118; BPp65ADZp (attP40); BPZpGAL4DBD (attP2) | Hampel et al., 2015 | Empty-SS |
| *D. melanogaster*: w1118; VT043699-p65ADZp (attP40) /CyO::Tb-RFP; 11A07-ZpGdbd (attP2) | Schretter et al., 2020 | SS36551<br>Available via https://www.janelia.org/split-GAL4. |
| *D. melanogaster*: w1118; VT064565-p65ADZp (attP40); VT043699-ZpGAL4DBD (attP2) | Schretter et al., 2020 | SS36564<br>Available via https://www.janelia.org/split-GAL4. |
| *D. melanogaster*: w1118; 35C10-p65ADZp (JK73A), 71A09-ZpGdbd (attP2) | Schretter et al., 2020 | SS56987<br>Available via https://www.janelia.org/split-GAL4. |
| *D. melanogaster*: w1118; 26A03-p65ADZp (attP40); 54A05-ZpGdbd (attP2) | Wu, Nern et al., 2016 | OL0046B<br>Available via https://www.janelia.org/split-GAL4. |
| *D. melanogaster*: w1118; 56H10-p65ADZp (attP40); 22E10-ZpGdbd (attP2) | Turner-Evans et al., 2020 | SS00238<br>Available via https://www.janelia.org/split-GAL4. |
| *D. melanogaster*: w1118; 16E11-p65ADZp (attP40); 21C11-ZpGdbd (attP2) | Namiki et al., 2022 | SS03500<br>Available via https://www.janelia.org/split-GAL4. |
| *D. melanogaster*: w1118; 38G07-p65ADZp (attP40); 20H05- ZpGdbd (attP2) | Meissner et al., 2024 | SS00030<br>Available via https://flylight-raw.janelia.org. |
| *D. melanogaster*: w1118; 38B06-p65ADZp (attP40); 26C06- ZpGdbd (attP2) | Meissner et al., 2024 | SS00168<br>Available via https://flylight-raw.janelia.org. |
| *D. melanogaster*: w1118; 52B10-p65ADZp (attP40); 20C08- ZpGdbd (attP2) | Meissner et al., 2024 | SS00217<br>Available via https://flylight-raw.janelia.org. |
| *D. melanogaster*: w1118; 81D05-p65ADZp (attP40); 89G01- ZpGdbd (attP2) | Meissner et al., 2024 | SS00277<br>Available via https://flylight-raw.janelia.org. |
| *D. melanogaster*: w1118; 16D01-p65ADZp (attP40); 22G07-ZpGdbd (attP2) | Meissner et al., 2024 | SS00006<br>Available via https://flylight-raw.janelia.org. |
| *D. melanogaster*: w1118; 56H10-p65ADZp (attP40); 32A11- ZpGdbd (attP2) | Meissner et al., 2024 | SS00243<br>Available via https://flylight-raw.janelia.org. |
| *D. melanogaster*: w1118; 71G01-GAL4 (attP40) | Pfeiffer et al., 2008<br>Jenett et al., 2012 | 71G01<br>Available via https://flweb.janelia.org. |
| *D. melanogaster*: w1118; 72C11-GAL4 (attP40) | Pfeiffer et al., 2008<br>Jenett et al., 2012 | 72C11<br>Available via https://flweb.janelia.org. |
| *D. melanogaster*: w1118; 20A02-GAL4 (attP40) | Pfeiffer et al., 2008<br>Jenett et al., 2012 | 20A02<br>Available via https://flweb.janelia.org. |
| *D. melanogaster*: w1118; 65F12-GAL4 (attP40) | Pfeiffer et al., 2008<br>Jenett et al., 2012 | 65F12<br>Available via https://flweb.janelia.org. |
| *D. melanogaster*: w1118; 91B01-GAL4 (attP40) | Pfeiffer et al., 2008<br>Jenett et al., 2012 | 91B01<br>Available via https://flweb.janelia.org. |

## Acknowledgments

We would like to thank the Janelia Experimental Technology group for design collaboration and fabrication assistance with the Fly Disco hardware including in particular Roger Rodgers and Jeff Talbot for bubble cartridge development and fabrication, and Steve Sawtelle for the LED board and controller development and fabrication. The Project Technical Resources at Janelia including Austin Edwards, Nan Chen and Mary Lay for keypoint annotations. Nan Chen, Austin Edwards, Rebecca Arruda, Claire Managan, and Kaitlyn Boone for the data collection over the years as we developed the Fly Disco and conducted various screens. The Invertebrate Shared Resource group for the production of innumerable flies for our experiments. Gerry Rubin for funding the replication and documentation of the Fly Disco hardware. William Dickson for BIAS software development. Allen Lee for software development in previous versions of the analysis pipeline. Igor Siwanowicz for the rendering of the fly body model.

